# Pollen Partners: The Symbiotic Microbes of *Pinus radiata* Pollen

**DOI:** 10.1101/2024.07.31.606094

**Authors:** C Armstrong, S Ganasamurthy, K Walker, C Mercier, S Wakelin

**Affiliations:** Scion, Christchurch 8011, New Zealand; Scion, Rotorua 3010, New Zealand

## Abstract

Pollen, a crucial source of nutrients and energy for pollinators. It also provides a unique habitat for ecological microbiota. Previous research on the microbiome of pollen has largely focussed on angiosperm systems, with limited research into coniferous gymnosperms. This study characterises the pollen microbiome associated with one of the world’s most widely grown tree species, *Pinus radiata*. Trees were sampled from locations across Canterbury, New Zealand, with repeated collections in 2020 and 2021. Metabolomic analysis revealed the main compounds present on *P. radiata* pollen to be amino acids (principally proline), and carbohydrates (fructose, glucose, and sucrose). Although phenolic compounds such as ρ-coumaric acid and catechin, and terpenoids such as dehydroabietic acid, were present at low concentrations, their strong bioactive natures mean they may be important in filtering of microbiome communities on pollen. *Pinus radiata* pollen was found to host a microbiome dominated by fungi; this directly contrasts with those for many angiosperm species. Geographic range and sampling years were evaluated as secondary drivers of microbiome assembly. Neither sampling location nor annual variation had a significant impact on the fungal component of the pine pollen microbiome, which was remarkably stable/conserved among samples. However, some bacterial taxa exhibited sensitivity to geographic distances and yearly variations, suggesting a secondary role for some. A core microbiome was identified in *P. radiata* pollen, characterized by a consistent presence of specific fungal and bacterial taxa across samples. While the dominant phyla, Proteobacteria and Ascomycota, align with findings from other pollen microbiome studies, unique core members were unidentified at genus level. This tree species-specific microbiome assembly emphasizes the crucial role of the host plant in shaping the pollen microbiome. These findings contribute to a deeper understanding of pollen microbiomes in gymnosperms, shedding light on the need to look further at their ecological and functional roles.

## Introduction

It is well established that plants harbour a wide diversity of microorganisms in the phyllosphere and other above- and below-ground tissues (Vorholt, 2012). Until recently, however, the microbiome associated with plant reproductive organs has been largely overlooked. Yet emerging research is showing that pollen, for example, can be host to diverse communities of bacteria, fungi, viruses, and other microbiota (Manirajan *et al*., 2016). The discovery that pollen has a microbiome has implications not only for individual plants and their reproductive success (Zasloff, 2017), but also has complex, ecosystem-level outcomes (Wardle *et al*., 2004).

Studies on the microbiome of pollen have largely focussed on angiosperm systems (i.e. flowering plants), which has been shown to impact ecosystems via influences on pollinator ecology (Cullen *et al*; 2021). For example, it has been posited that global decline in bees is linked to the use of agricultural chemicals that result in dysbiosis of the normal pollen microbiome that pollinators require and/or have an immune defence against (Zasloff, 2017). As pollen is a major dietary resource for bee larvae, the microbial community on pollen is a major factor influencing the health and fitness of these pollinator species (Dharampal *et al.,* 2019). Additionally, links between pollen microbiome and human allergenicity are evident (Manirajan *et al.,* 2019). Bacterial toxins can add to the load of allergens on pollen with outcomes on human health (Manirajan *et al.,* 2022; Obersteiner *et al.,* 2016). Thus, from complex and idiosyncratic impacts on structure and functioning of ecosystems through to public health, the implications of the pollen microbiome are profoundly impactful yet vastly understudied.

Pollen represents a vital transport shuttle for movement of microorganisms onto, across tissues, and among plants. It facilitates dispersal of microorganisms within macro-ecosystems and transmission across larger geographic ranges (Sessitsch *et al.,* 2023). The pollen microbiome is, of course, ideally situated for colonisation of the ovule and seed after pollination (Argarwal and Sinclair, 1997). This supplies a direct path for vertical (paternal) microbiome inheritance and coevolution with the host (Nelson, 2018; Wassermann *et al.,* 2022; Vannier *et al.,* 2018). Similarly, this process provides a pathway for pathogens or other symbionts to enter and impact plant organs grown for productive purposes such as nuts and fruits (Donati *et al.,* 2018; Cellini *et al.,* 2019). Pollen can contain significant amounts of mono- and oligo-saccharides (sugars), nitrogen, protein, micronutrients such as K, S, Cu, Fe, Zn, and other metabolites (Conti *et al.,* 2016; Filipiak, 2016). During transport in the environment, this may provide resources that sustain epiphytic microbial cells. Conversely, other microorganisms actively parasitise pollen, utilising the material as a resource base *per se*, and as a source of transported inoculum for decomposition of litter and other debris within the wider ecosystem (Hutchinson and Barron, 1997).

Forest plantations are an important source of ecosystem services such as wood and fibre. The fast growth rate of plantation forest trees enables them to supply of 35% of world’s wood supply while comprising only 5% of global forest area (FAO, 2010). Increases in demand for wood and other services, including biofuel, food, and other forest bioproducts can come either at the expense of large natural ecosystems or more intensively managed productive forests (FAO *et al.,* 2012). In order to grow the global bioeconomy, increased utilisation of highly productive planted-forests may not only provide raw carbon-neutral resources, but avoid biodiversity loss and other ecosystem impacts associated with sourcing material from natural forest systems.

*Pinus* spp. are extensively used in planted forests globally (FAO, 2006). When plantation forests are grown as monocultures, these areas of intensive *Pinus* spp. Provide massive pulses of pollen into the ecosystem annually; indeed, this as key phenological event occurring in forested ecosystems (e.g. López-Orozco et al., 2023; Schermer et al., 2020). In New Zealand, for example, approximately 90% of the entire planted forests comprise *Pinus radiata* (FOA, 2016). This single species covers an extent of 1.6 m Ha of land area and, based on the estimates of Fielding (1960), may release ∼ 300 million kg of pollen each spring (∼270 kg per ha, depending on site, environment, silvicultural regime, and tree genetics). This comprises a massive input of carbohydrate, nitrogen, other nutrients, into the ecosystem (Hutchinson and Barron, 1997; Greenfield 1996). It might also facilitate a massive input and exchange of microbiomes from the tree to the environment. From a public health perspective, this is certainly nothing to sneeze at. However, the wider ecological significance of such a heavy pulse of pollen and its microbiome into the environment have yet to be investigated.

The purpose of this study is to expand our knowledge of the occurrence of pollen microbiome of coniferous, gymnosperm tree species. Conifers are important globally, dominating the composition of boreal forests which comprise 24% of total global tree cover alone. Conifers are also important in temperate forest systems that span Europe, Asia, the Americas, Africa, and Australasia. With a few exceptions, most conifers are needle-leaved evergreen trees; these make up ∼38% of the trees present globally (Ma et al., 2003). Given the focus of (already limited) pollen microbiome studies towards angiosperm spp., this helps addresses a huge gap in our knowledge *per se*. However, the primary focus on *Pinus radiata* is because it provides a useful model system for tree-microbiome research (Addison et al., 2003a, 2003b, 2024). In addition to describing pine pollen microbiome, we aimed to determine how much of the community is variable versus stable/conserved by sampling trees across a wide geographic range and across two consecutive seasons (years). Finally, given the focus on exploration of a new microbiome habitat, we used a non-targeted metabolomics approach to determine the range of potential metabolites on the surface of *Pinus radiata* pine pollen that may provide resources, or inhibitory molecules, that may support microbial growth or provide a filter for microbiome community composition.

## Methods and Materials

### Sample collection

Twenty-four *Pinus radiata* trees were randomly selected across the Canterbury region in New Zealand (Figure 1). This region provides locations ranging from coastal areas, planted and natural forests, recreational parks, agricultural land, rivers, and the foothills of the Southern Alps. The selected trees were mature, exhibited good health without observable indications of stress or disease, and had accessible male cones (microsporangiate strobili) at the beginning of spring (September) when they begin to open and release pollen. The male cones of *P. radiata* are often referred to as catkins (Payn *et al.,* 2017), however these differ to catkins typically associated with angiosperm trees such as birches, alders, willows and others. Unless stated otherwise, we use the term catkin in respect to *Pinus-*type microsporangiate strobili. Pollen from the same trees was sampled in 2020 and 2021.

**Figure 1:**
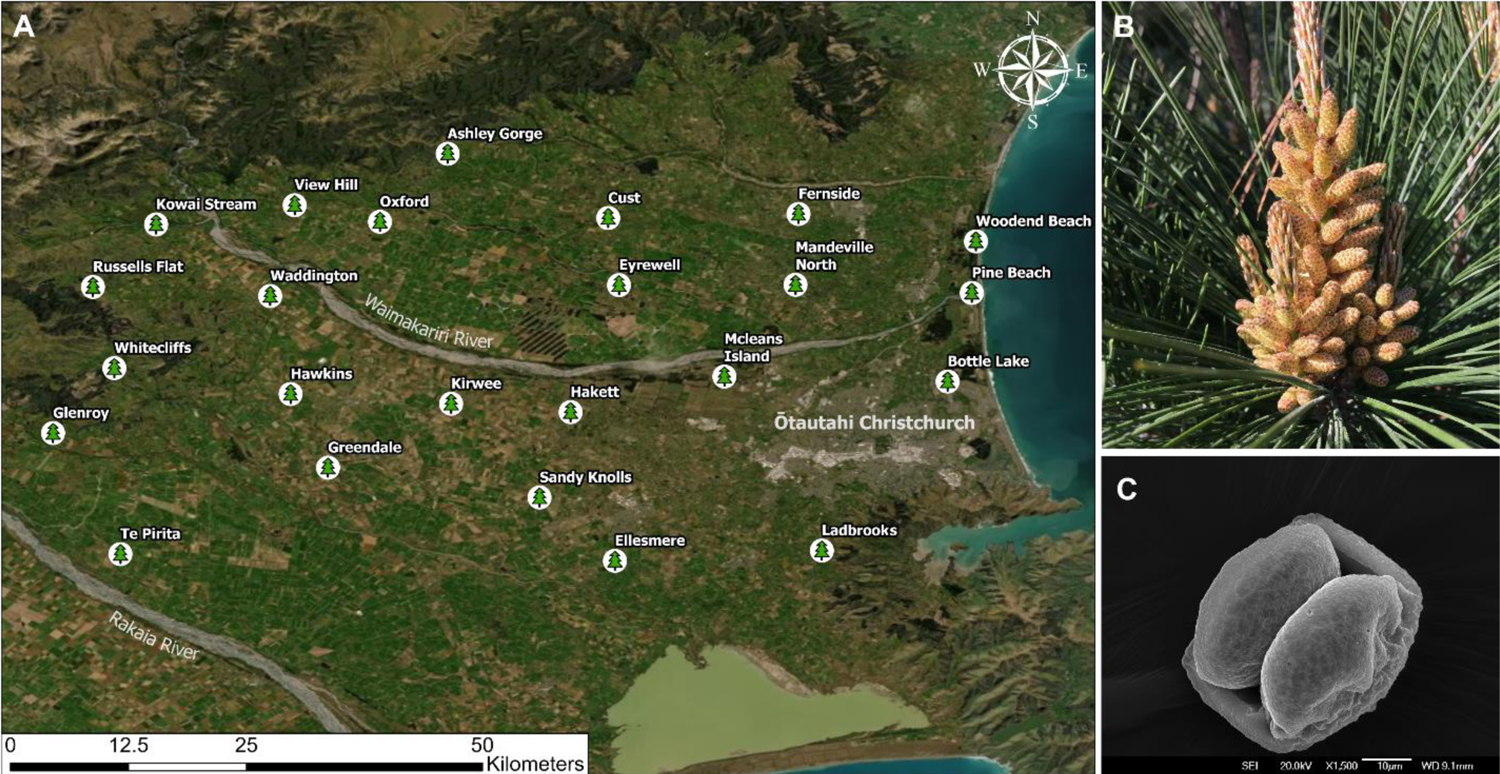
(A) Locations of the 24 *Pinus radiata* trees from which pollen was sampled from. Green pine symbols denote the locations of the trees and the name of the location is next to this symbol. The map is a satellite image of Canterbury based on *Google Earth* imagery for September 2021. (B) A cluster of *P. radiata* catkins (microsporangiate strobili) in September 2021, showing the catkins at pollen-release stage. (C) Scanning electron microscope (SEM) image of a *P. radiata* pollen grain; scale bar 10 µm.

For each tree sampling, a group of catkins clustered along a stem were excised off the tree using sterilised scissors. These were placed in a sterile tube and pollen was removed from the catkins by gentle shaking. The tubes were chilled until returned to the laboratory where they were stored in a −80°C freezer until all samples were collected.

Surface features and microbiome of pollen from selected trees was examined under a scanning electron microscope (SEM). Fresh pollen grains were initially sputter-coated with palladium (Emitech K975X coater) before being placed in a JEOL JSM IT-300 variable pressure SEM. Microscopy was conducted at the University of Canterbury Electron Microscopy Centre (Earth and Environmental Sciences Department). Similar to other Pinaceae species, *P. radiata* has bisaccate pollen, in which two air sacs protrude from the pollen wall. These sacs facilitate long distance wind dispersal (Schwendemann *et al.,* 2007).

The metabolome resource available for microorganisms on the surface of *P. radiata* pollen was characterised. Specimen *P. radiata* trees, growing adjacent to a commercial plantation forest at McLeans island (Fig 1A) were sampled in 2023. At the site, 12 samples of pollen were collected from different catkin structures. Each sample was gently rotated in 3:3:2 ACN:IPA:H2O (acetonitrile, isopropanol, water) solution to extract polar and non-polar metabolites off the pollen surface. The ACN:IPA:H2O extractant was processed by West Coast Metabolomics (California, USA) and compounds were separated and identified using gas chromatography - time-of-flight - mass spectrometry (GC-TOF-MS). Data was provided as normalised peak heights to represent the relative semi-quantification of each compound.

### DNA Extraction, library preparation and amplicon sequencing

Genomic DNA was extracted from 50-100 mg samples pollen using the DNeasy PowerSoil DNA Extraction Kit (Qiagen, Germany) following the manufacturer’s recommended instructions. DNA was processed for bacterial and fungal sequencing, targeting the respective 16S rRNA and ITS housekeeping gene regions. The primers used to amplify these genes were based on protocols from the Earth Microbiome Project (Caporaso *et al.,* 2012). The 16S rRNA primers were 515F (5’-GTGYCAGCMGCCGCGGTAA-3’) and 806R (5’-GGACTACNVGGGTWTCTAAT-3’) (Parada *et al.,* 2016) and ITS amplification used primers ITS1f (5’-CTTGGTCATTTAGAGGAAGTAA-3’) (Ihrmark *et al.,* 2012) and ITS2 (5’-GCTGCGTTCTTCATCGATGC-3’) (White *et al.,* 1990). Primers were barcoded using the Golay 12-mer system, including Illumina adapters. Amplicons were cleaned using a magnetic bead PCR clean-up kit (Geneaid, Taiwan) and pooled to an equimolar mix. The 16S and ITS libraries were sequenced using Illumina MiSeq sequencing with 250 bp pair end (PE) read chemistry (16S) and 300 bp PE read chemistry (ITS); this was conducted at the Australian Genome Research Facility (AGRF).

### Bioinformatics and data processing

Raw sequencing data was processed using the DADA2 pipeline outlined for 16S and ITS tag sequences (Callahan *et al*., 2016). Chimeric sequences were removed using the removeBimeraDenovo tool provided by DADA2, and taxonomy was assigned to amplicon sequences variants (ASVs) using the assignTaxonomy tool and the RDP reference database for 16S (Cole *et al*., 2014) and the UNITE database for ITS (Kõljalg *et al*., 2005). Bacterial and fungal ASV count and taxonomy tables were filtered to remove unassigned sequences, singletons, and samples with less than 150 reads. Additional filtering to remove chloroplast sequences was performed for the 16S rRNA gene dataset. A phyloseq object was created in R by merging the ASV count table, taxonomy table, and metadata; this was the base for much of the subsequent data visualisation and statistical analysis.

### Statistical analysis and Graphs

Statistical analyses were performed in R version 4.0.5 (R Core Team, 2021) and PRIMER/PERMANOVA+ (PrimerE Ltd.). Alpha diversity measures were calculated via the ‘phyloseq’ package (McMurdie and Holmes, 2013), boxplots were created using GraphPad Prism version 9.4.1 (GraphPad Software, USA) and p values were computed using a Mann-Whitney test (between two groups). Rarefaction curves were produced with the vegan package (Oksanen *et al*., 2022).

To identify factors potentially influencing pollen microbiome composition, PERMANOVA modelling (Anderson, 2001) was conducted with sampling year and cardinal location (see later) as independent, fixed variables. These were conducted in PRIMER (Anderson, 2001) using Bray-Curtis calculated distances. Non-metric MDS plots of these distances were then generated in R. The metacoder package (Foster *et al*., 2017) was used to compute the hierarchical distribution of taxonomic groups in terms of abundance. The clustered heatmap was computed using the pheatmap (Kolde, 2012) package in R. The core microbiome plots were generated using the UpsetR (Conway *et al*., 2017) and microbiome packages (Lahti *et al*., 2017). All plots were generated using ggplot2 (Wickham, 2016) unless otherwise stated.

In the initial testing of main effects, the influence of geographic location on microbiome composition was tested by grouping the samples together based on splitting the sample area into cardinal location quadrants from an approximate centroid. This testing allowed for a ‘cardinal location’ term to be included in the model alongside sampling year. However, this was exploratory in nature and did not provide a consistent means of investigating this variable. In order to circumvent this and identify if microbiome changes with location, geographic distances were calculated and correlated against class-level calculated phylogenetic distance matrices (Bray-Curtis) using Spearman’s-rank correlation. This initial testing involved testing of relationship between geographic distance and all bacterial taxa combined. Geographic distances were computed using the Geographic Distance Matrix generator (Ersts, 2023) based on sample latitude and longitude input datum.

To partition which individual groups within the community exhibited distance-decay relationships, secondary testing was conducted using the BIO-ENV routine (Clarke 1993; Clarke and Ainsworth 1993). In this approach, distance matrices (Bray-Curtis) were created individual and combinations of class-group of bacteria and rank-correlations among these and geographic distance was calculated. A permutation-based approach was then used to test for significance of the effects.

Metabolome data were averaged across the 12 samples to obtain a single, representative, *P. radiata* pollen metabolome. The data was classified into superclass groups and the relative quantifications of the compounds were standardised as a percentage of total compounds per sample. Note that this is purely descriptive analysis. No formal testing of location or other effects were tested, rather a typical ‘fingerprint’ of the pollen metabolome generated.

## Results

### Bacterial and fungal microbiome richness

Alpha diversity of individual *P. radiata* pollen microbiomes were calculated as observed richness (Kim *et al.,* 2017). The data is shown in Figure 2, with richness of prokaryotes and fungi for years 2020 and 2021. The richness of fungal taxa on pollen was much greater than for prokaryotes, averaging over 200 (2020=201; 2021=212) ASV’s per sample compared to the just over 40 (2020=41; 2021=44) for bacteria. For both fungi and bacteria, the richness of microbiomes present on pollen was consistent across the sampling years (Wilcoxon-test; bacteria p=0.81, fungi p=0.96).

**Figure 2:**
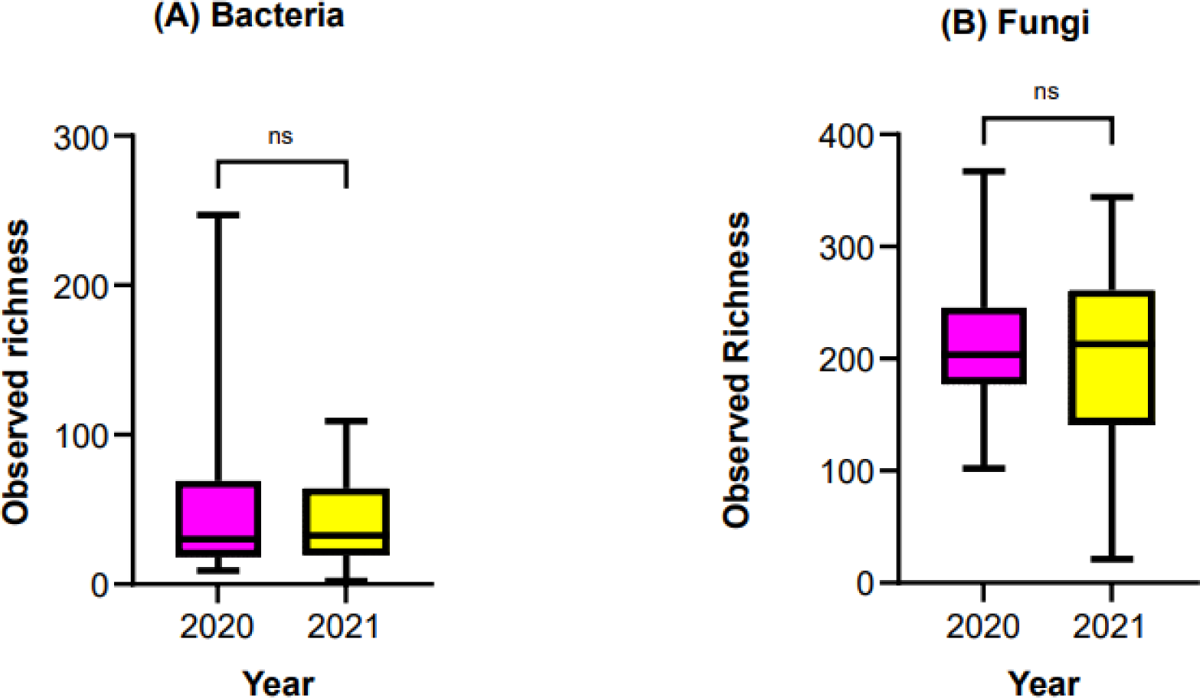
Observed richness of bacteria (left) and fungal (right) taxa present on *Pinus radiata* pollen (24 sampling sites) collected over two years.

As total microbiome richness per sample was relatively low (i.e. tens- to hundreds of taxa per sample), we expected the MiSeq sequencing runs to capture the full extent of taxa present. Indeed, doing so can be important for increasing the accuracy of alpha-diversity estimates as well as down-stream investigation of community composition. To confirm this, rarefaction curves were generated; these are given in Figure S1. After clean-up and processing, the majority of 16S sequencing (28/33 samples) had <4,000 tag sequence reads. This was due to dominance of plant chloroplast sequences in the libraries alongside relatively low bacterial richness (Fig 2 A). Conversely, as more fungal sequences passed through QC, sequencing depth was greater with all samples having from 16,000 and 150,000 useful read for ASV and taxonomic assignment.

### Composition of the *Pinus radiata* pollen microbiome and defining the main taxa present

The microbiome of *P. radiata* pollen is dominated by fungi, having 3.7-fold greater ASV counts in comparison to bacteria. These findings are in line with measures of observed richness (i.e. Fig. 2). The fungal community was dominated by Dothidiomycetes (Ascomycetes), and Tremellomycetes (Basidiomycete). A few other sub-phyla including Mucoromyceta, Mortierellomycota, and Eurotiomycetes were also present, but at lower abundances (Figure 3).

**Figure 3:**
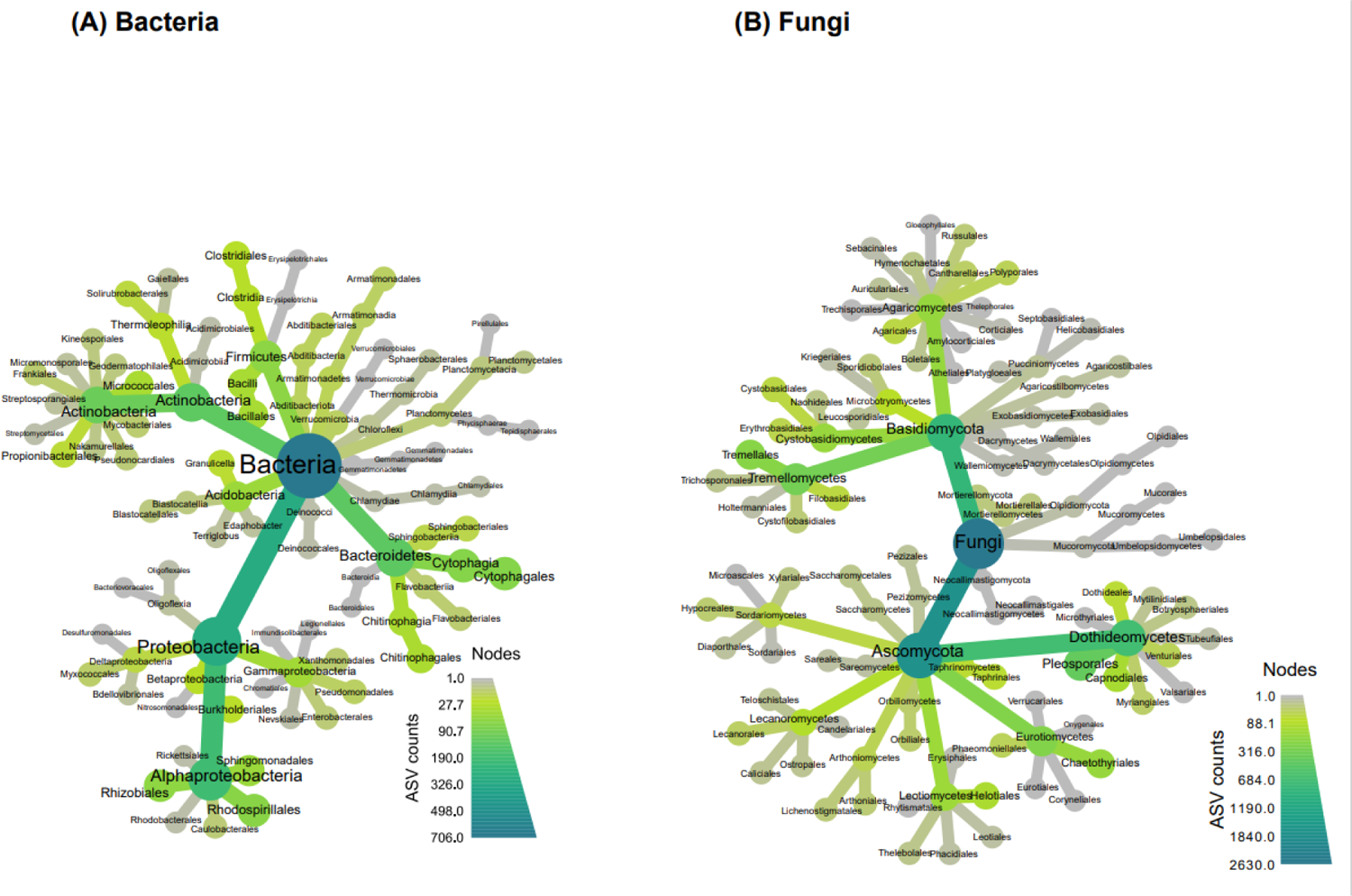
Phylogenetically hierarchical distribution (‘metacoder plots’) for bacteria (left) and fungal (right) microbiomes present on *Pinus radiata* pollen. Darker gradient fill indicates increase in ASV count of taxa on branches and leaves. Plots combine data for sites and years to indicate the overall spectrum of taxa found. Size of nodes indicate ASV counts.

The bacterial taxa present in the pollen microbiome were more evenly distributed across the phyla and classes present than was observed for the fungal microbiome. However, trends were still evident. Within the Proteobacteria, Alpha-proteobacteria were numerically important. Similarly, the taxa present in the phylum Actinobacteria were concentrated within the class Actinobacteria (Figure 3). Thus, although a greater range of phyla was present compared with fungi, the diversity within each phyla was lower.

Data was retained at class-level aggregation for inspection of dominance and variation of taxa among samples providing some context towards the observations in Figure 3. These are presented in Figure 4A for bacteria, and Figure 4B for fungi.

**Figure 4:**
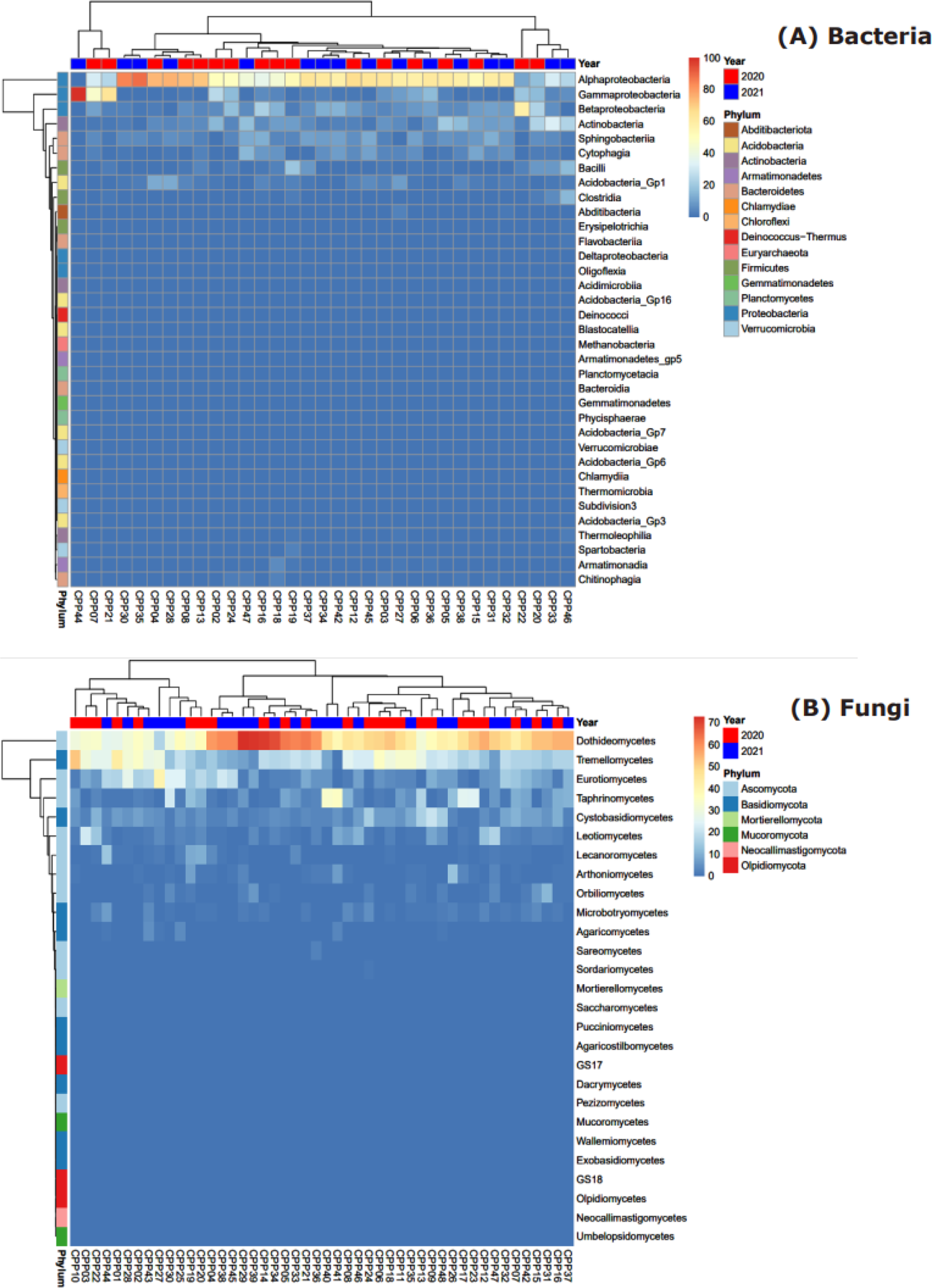
Relative abundance (%) heatmaps of bacteria (top) and fungi (bottom) *Pinus radiata* pollen microbiomes at class level. The x-axis denotes individual sample locations; coloured tiles on the top of each heatmap indicate sampling years and coloured tiles on the left indicate corresponding phylum

The bacterial community of the *P. radiata* pollen microbiome is dominated by Proteobacteria and, in, particular Alphaproteobacteria (Fig 6A). The only other phyla to have numerically meaningful abundances at class level were Actinobacteria, Bacteroidetes (Sphingobacteriia and Cytophagia), Firmicutes (Bacilli and Clostridia), and Acidobacteria (GP_1). Each of these groups exhibited sample-specific variation. For example, Betaproteobacteria was the most abundant in the sample collected from the Glenroy 2020 collection (relative abundance > 50%), but was either absent or was only present in the range of 0.9 to 23.3% for all other samples. In contrast, Alphaproteobacteria was not only the most abundant amongst the samples (ranging in relative abundance from 11 to 88.3%), but was also present in nearly all samples. The only exception was a sample collected from Russel Flats in 2021; this was comprised entirely of Gammaproteobacteria.

The fungal community was dominated by Ascomycota and Basidiomycota taxa (Fig 6B). Dothiodeomycetes were the most abundant group across samples, ranging in relative abundance from 16.9% to 72.8% of the fungi present, and next were Tramellomycetes (0.9 to 52.3% relative abundance). As for the bacteria, individual pollen samples exhibited anomalously high abundances of some fungal groups. For example, samples collected from Ashley Gorge in 2021 and View Hill in 2021 had 33.9% and 35.3% of Taphrinomycetes, respectively, but other samples had as little as 0.2% or up to 25.6%. A similar trend was also observed across classes Eurotiomycetes and Cystobasidiomycetes.

### The pine pollen metabolome

The prominent metabolites present on the surface of *P. radiata* pollen are amino acids and organic oxygen compounds (primarily carbohydrates and alcohols). These comprised ∼81% of the pine pollen metabolome (Fig 5). At the compound level, proline was by far the most dominant metabolite, present at 17.8%. Other amino acids with notable abundances were alanine, γ-aminobutyric acid (GABA), valine, and oxoproline (1.4 - 3.8%). Of the organic oxygen compounds, pinitol (a cyclic polyol that’s been found to be associated with other pine trees) and glycerol were both of high abundance, at ∼5%. Fructose, glucose and sucrose were the three most prominent sugars at 2-3%. The most prominent lipids present were two prenol lipds: dehydroabietic acid (3.19%) and abietic acid (1.8%). Several phenolic compounds and flavonoids such as ρ-coumaric acid and catechin were also found, although at low concentrations (<0.1% per compound).

**Figure 5:**
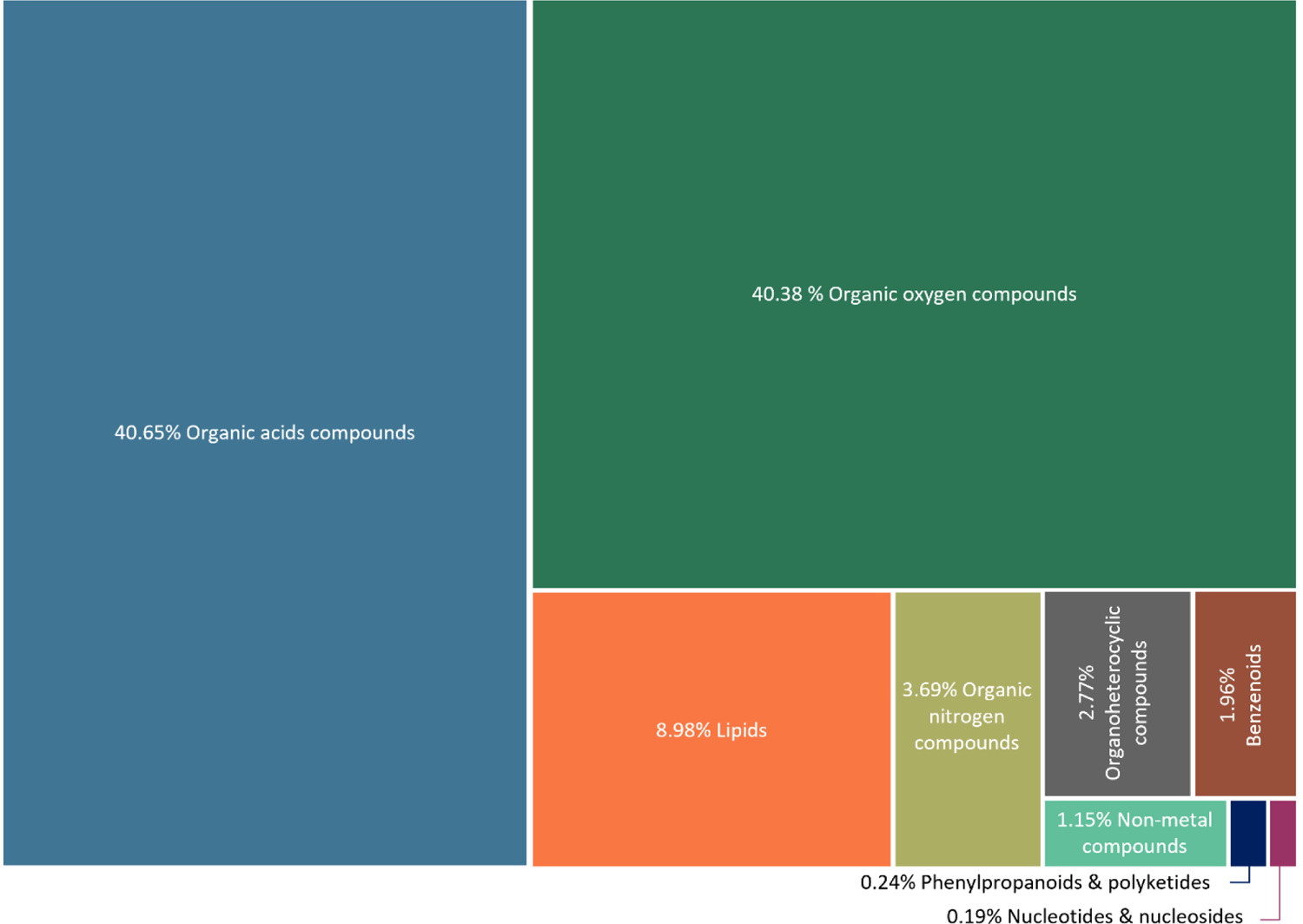
Primary metabolite compounds found on the surface of *Pinus radiata* pollen. Compounds are grouped at superclass level and the relative abundance of the metabolites are standardised as a percentage of total compounds from 12 pollen samples.

### Does geographic distance or annual variation influence pollen microbiome composition?

The influence of cardinal location or year on pollen microbiome composition was tested using PERMANOVA; summary results are presented in Table 1. The pollen fungal community composition was stable over the two sample time points (p=0.132) and with cardinal location (p=0.601). No interaction effect was detected between the two factors (Table 1). Similarly, in the nMDS ordination, no structure or separation was evident related to cardinal location nor year of collection (Fig. 6). Thus, the pollen associated fungal community was stable both over time of collection and with cardinal location grouping.

**Figure 6:**
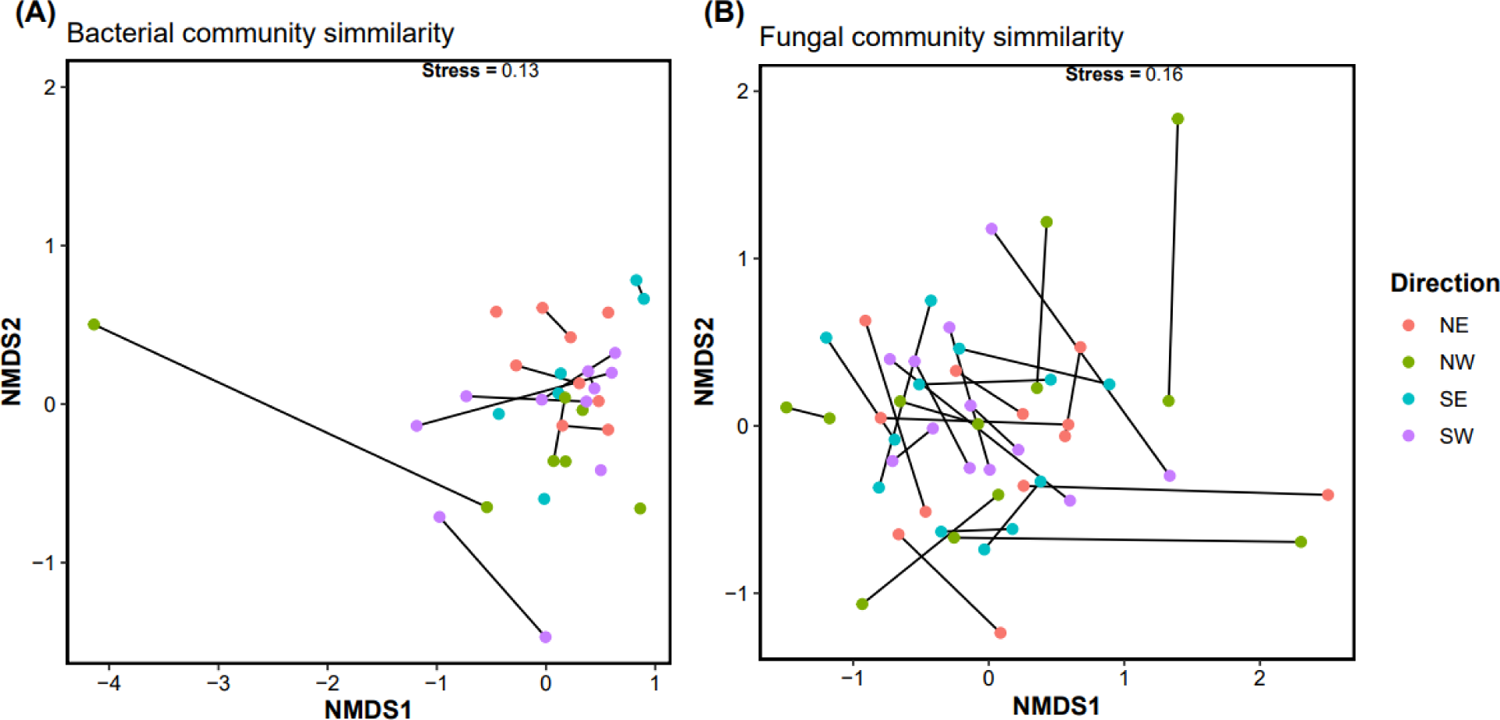
nMDS ordination plots showing similarity among samples of pollen in bacteria. (A) and fungal (B) community composition by sampling year (black line) and cardinal direction location (coloured dot). Samples from the same location are connected by year of sampling. Note that some pollen samples had no bacterial microbiome in one of the sampling years; these locations therefore appear as single points.

**Table 1:**
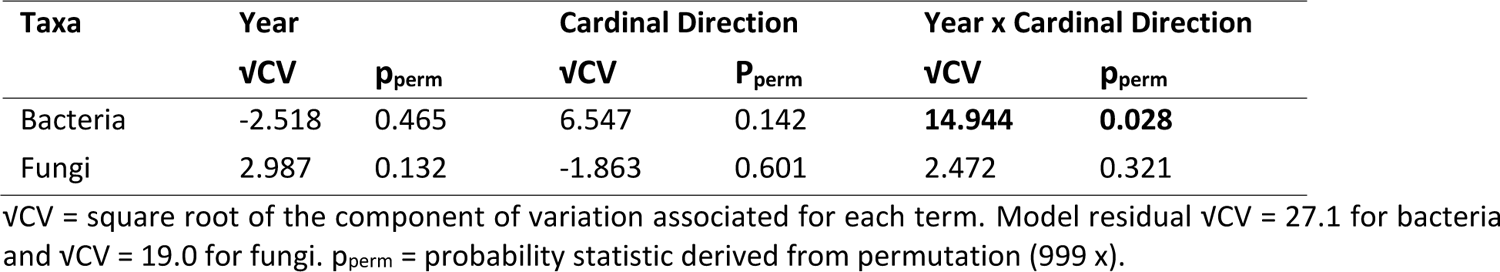
Summary PERMANOVA results table testing effects of year, cardinal location and their interaction on the bacterial and fungal microbiome composition of *Pinus radiata* pollen.

Similarly, bacterial community showed no variation directly related to sampling year (p=0.465), nor sampling location (cardinal direction p=0.142) (Table 1). However, an interaction term of year x cardinal direction was evident, and the variation component associated with this was strong (√CV=14.944; p=0.028). Exploration of this effect (pair-wise tests) suggested an influence of sampling across the SW v NW quadrants in relation to microbiome composition in 2020 (p=0.059) and in 2021 (p=0.074).

The grouping of samples into cardinal quadrants was used to explore the broad influence of location in the PERMANOVA based testing and apportion this variance (if any) relative to year and test for interaction effects. However, this grouping can be somewhat arbitrary and may mask secondary underlying relationship among biologic or abiotic factors associated with microbiome assembly over this sampling area. For example, samples collected at points close to the geographic centroid of the collection range could be separated into NE, SE, NW, and SW groups, yet would be effectively adjacent in absolute distance. The opposite would be true at the edge of the sampling area. As such, further analysis was conducted based on calculated pair-wise distances among sample locations to provide robust determination of the relationships between change in biological communities with geographic range.

Towards this, Mantel-based testing was used to formally test for evidence of association (rank-correlation) between the geographic and biological-based distance matrices. For the fungal community, weak correlation (ρ=0.089) was evident. However, evidence for association between changes in bacterial community and sampling range was stronger (ρ=0.227). Given this, only bacterial community associations were further explored.

Using BIO-ENV matching, changes in bacterial community that were associated with spatial variation were tested. Individual taxa exhibiting strongest distance-dependent relationships were Clostridia (ρ=0.225), candidate division WPS-1 (ρ=0.217), Sphingobacteriia (ρ=0.213), and a non-classified Proteobacteria class (ncC_Proteobacteria; ρ=0.184). Higher correlations were present when bacterial groups were combined; the highest included either Sphingobacteriia or Clostridia in combinations with the ncC_Proteobacteira, Alphaproteobacteria, WPS-1 group (as before), or other taxa such as Oligoflexia or Acidobacteria. Regardless, Sphingobacteriia and Clostridia were both important but could be included both exchanged for each other in tests that bacterial groups. The combination of taxa in which highest correlation with geographic distance was found included Sphingobacteriia, non-classified Proteobacteria, Clostridia, Oligoflexia and either Acidimicrobiia or Deinococci (ρ=0.346, p=0.005). Relationships between change in taxa abundances and geographic sampling are given in Fig. S2.

### The core microbiome of *Pinus radiata* pollen

Core genera were defined as those having a minimum 50% prevalence and 0.01 detection threshold across samples (Graystock et al., 2017; Kardas et al., 2023). Detection threshold is the minimum relative abundance (i.e. 1%) value at which groups are considered present in the community while prevalence of 50% indicates that groups must be present in at least 50% of the samples to be considered a core member. The core microbiome conserved across all samples (locations and years) are summarised in Table 2. Overall, fungi had twice as many core genera compared to bacteria with relative abundance being genus-specific. An UpSet plot showing core genera between and across years and highlighting subtle sampling time-based influences are shown in Figure S3.

**Table 2:** The core bacterial and fungal genera, prevalent across at least 50% of the samples at a minimum of 0.01 detection threshold. The relative abundance ranges from the lowest abundance in a sample to the highest abundance in a sample.

| Bacteria | Relative abundance % | Phylogenetic lineage |
| --- | --- | --- |
| Robbsia | 0.85-23.26% | Proteobacteria; Betaproteobacteria; Burkholderiales; Burkholderiaceae |
| Sphingomonas | 2.22-23.23% | Proteobacteria; Alphaproteobacteria; Sphingomonadales; Sphingomonadaceae |
| Hymenobacter | 0.45-12.55% | Bacteroidota; Cytophagia; Cytophagales; Hymenobacteraceae |
| Acidisoma | 0.63-5.13% | Proteobacteria; Alphaproteobacteria; Rhodospirillales; Acetobacteraceae |
| Methylobacterium | 0.3-5.79% | Proteobacteria; Alphaproteobacteria; Hyphomicrobiales; Methylobacteriaceae |
| Fungi | Relative abundance % | Phylogenetic lineage |
| Microsphaeropsis | 0.1-46.24% | Ascomycota; Dothideomycetes; Pleosporales; Didymosphaeriaceae |
| Perusta | 0.12-44.26% | Ascomycota; Pezizomycotina; Dothideomycetes; Dothideomycetidae; Dothideomycetidae incertae sedis |
| Hormonema | 0.12-34.01% | Ascomycota; Pezizomycotina; Dothideomycetes; Dothideomycetidae; Dothideales; Dothioraceae |
| Taphrina | 0.14-30.98% | Ascomycota; Taphrinomycotina; Taphrinomycetes; Taphrinales; Taphrinaceae |
| Gelidatrema | 0.02-24.76% | Basidiomycota; Agaricomycotina; Tremellomycetes; Tremellales; Phaeotremellaceae |
| Naganishia | 0.06-10.28% | Basidiomycota; Agaricomycotina; Tremellomycetes; Filobasidiales; Filobasidiaceae |
| Vishniacozyma | 0.13-10.32% | Basidiomycota; Agaricomycotina; Tremellomycetes; Tremellales; Bulleribasidiaceae |
| Genolevuria | 0.08-8.66% | Basidiomycota; Agaricomycotina; Tremellomycetes; Tremellales; Bulleraceae |
| Epicoccum | 0.21-8.36% | Ascomycota; Dothideomycetes; Pleosporales; Didymellaceae |
| Symmetrospora | 0.17-4.88% | Basidiomycota; Pucciniomycotina; Cystobasidiomycetes; Cystobasidiomycetes incertae sedis; Symmetrosporaceae |

## Discussion

### Pine pollen hosts a fungal dominated microbiome

Our results demonstrated that *Pinus radiata* pollen does host a microbiome, and this microbiome is an order-of-magnitude richer in fungal taxa compared with bacteria. This finding differs to reports on various angiosperm species where the pollen is richest in bacterial taxa. Manirajan *et al*. (2018), for example, investigated pollen grains from 8 species: birch/*Betula*, canola/*Brassica*, rye/*Secale*, autumn crocus/*Colchicum*, common hazel/*Corylus*, blackthorn/*Prunus*, common mugwort/*Artemisia*, and cherry plum/*Prunus*. Each individual plant species had a characteristic pollen microbiome and, among these, differences between wind and insect-pollenated species were then present. However, across all plant species and pollen traits examined, the bacteria and fungi had a similar number of OTUs and the bacteria was more diverse than the fungi. Similar findings were observed in the works of Obersteiner *et al*. 2016 when looking at the pollen of birch (*Betula*) and timothy grass (*Phleum*). Regardless, the finding of fungal dominance of the pine-pollen microbiome is important as this is fundamentally different to that of angiosperm species.

A unique attribute of pine pollen is the durable exterior coating (exine) made of sporopollenin. Sporopollenin is one of the most chemically inert biological polymers known and helps ensure pollen is chemically resistant and environmentally durable (Stanley and Linskens, 1974). Each pollen exine has a unique and complex surface topology and chemistry enabling specificity with stigmas of reproductively compatible flowers. Immediately below the exine is an ‘intine’ inner later. Intine is composed of cellulose and hemi-cellulose, pectin and other proteins, and callose, and supports and regulates the growth of the pollen tube during fertilization. In some respects, pollen should comprise a preferred habitat for fungal colonisation. For example, the hydrophobicity properties of the exine layer (also pectin within the pollen) can favour fungal groups possessing surface-active hydrophobin proteins, allowing for attachment onto hydrophobic and hydrophilic surfaces (Wösten *et al.,* 1994; Takahashi *et al.,* 2005), such as pollen. Furthermore, antimicrobial peptides are present on surfaces of some pollen (Doughty *et al.,* 1998). These are hypothesised to play a role in preventing self-pollination of angiosperms (Nasrallah, 2002), but may also constrain the diversity of bacteria present on pollen surfaces (Zasloff, 2017). These factors combined could initially be considered to facilitate colonisation of fungal groups over bacterial groups in this unique microhabitat. However, many of these are properties typical of all pollen – not just those from conifers. Furthermore, and as noted previously, most studies describe a greater richness in bacterial species on pollen. As such, additional or alternative explanations need to be considered. These could be, for example: (1) strong host-level filtering based on selective chemicals, such as peptides noted by Zasloff (2017) but likely including other groups; (2) the pollen microbiome is a reflection of the catkin microbiome or other microbiomes present on the pine which is, in itself, fungal dominated and contributes taxa to the pollen; (3) the environmental microbiome is a dominant contributor onto pollen and unique properties of conifer-forest systems resulting in fungal dominance are reflected in the assembly of microbiomes on pollen.

Pine pollen is a known useful source of nutrients such as carbohydrates, proteins, lipids, phenolics, and minerals, which are essential for the wider ecosystem microbiota (Graham *et al.,* 2006, Pawlik and Ficek, 2023). It is also used as nutrient supplements in human health (Cheng *et al.,* 2023). With this in mind, it is not surprising to find that pollen is a suitable habitat for growth of a multitude of bacteria and fungi. Whilst there has been considerable research on the chemical composition of pine pollen, the nutrients present vary depending on the pine species itself as well as environmental factors such as altitude and soil characteristics (Cheng *et al*., 2023). Moreover, the process of undertaking chemical analysis can be destructive and, as such, the final data set often includes intracellular nutrients / metabolome as well. Given this, it is not always evident what metabolites are surficial on pollen and available for the microbes inhabiting this habitat.

Our analysis found that the *P. radiata* pollen surface metabolome contained a high percentage of amino acids and sugars, both of which are important nutrient and energy resources for microbial growth. Of particular note was proline being, by far, the most abundant metabolite present. Other pine pollens also exhibit significantly high levels of proline, including *Pinus jeffreyi* (Jeffrey pine) and *Pinus contorta* (Lodgepole pine), (Axelrod *et al*., 2021). The accumulation of proline in plants has been observed as a reaction to environmental stresses, including exposure to UV radiation (Saradhi *et al*., 1995; Szabados & Savoure, 2010). As pollen is transported by the wind, it becomes exposed to UV radiation. The presence of proline may serve as a protective mechanism for the haploid genome within the pollen.

There were also several flavonoids and polyphenols present and these are known to have antimicrobial properties that shape microbial communities on plant phyllospheres. Specifically, ρ-coumaric acid was the most abundant phenolic compound, which is a known antibacterial agent (Ojha and Patil, 2019), the presence of compounds like this could result in few bacteria present on the pollen, however, this is just speculation. Dehydroabietic acid (terpenoid) is found in resin and woody material of pine species. It has a broad range of metabolic and biological activities, including antimicrobial (Hao *et al.,* 2022). While a major constituent of resins and associated biomass burning and commercial lumber processing mills (Lai *et al.,* 2015), dehydroabietic acid is rarely reported in pollen. A notable exception is Jeffrey pine pollen (Axelrod *et al*., 2021). In addition to potential implications for microbiome selection/filtering on pollen, the presence of dehydroabietic acid on pollen needs to be considered in terms of its interaction with hydroxyl radicals and subsequent influence on atmospheric chemistry (Axelrod *et al*., 2021).

### Geographic range and time are secondary influences of microbiome assembly

The environment is a natural and important source of microbiomes for plant tissues (Sessitsch *et al.,* 2023). Indeed, there is ongoing bi-directional exchange of the microbiome between plants and the environment such that the systems are, in effect, an intimately interconnected meta-holobiont (Vannier *et al.,* 2018). However, whilst this is apparent for tissues such leaves and roots (Sessitsch *et al.,* 2023), the connection between the pollen and environment isn’t as well defined. The work of Obersteiner *et al*. (2016) demonstrated that even on different plant species occurring in the same location, the microbiome is most likely originating from the host plant; the environment was a second-order driver, with plant host-specific factors being most important. Similarly, Manirajan *et al*. (2016) assessed the pollen microbiome of plant species situated in and across different geographic areas and found pollen microbiomes were primarily host-specific. These studies indicate that the pollen microbiome is primarily influenced by factors associated with the host plant.

We explored if different sampling locations influenced assembly of pine pollen microbiome. In this study, *P. radiata* was collected within a 2,500 km^2^ sampling area and included trees in general proximity to a wide range of land uses. We initially grouped samples to cardinal locations relative the centre point of sampling [as noted earlier, these groupings were subjectively assigned and only used for exploratory analysis]. Sampling location/environment had no direct influence on the bacterial nor fungal microbiome on *P. radiata* pollen. We further tested to see if geographic distances among sampling locations was related to microbiome assembly. For example, in our sampling region the eastern areas are coastal and have a drier and warmer climate than those close to the mountainous (further west). In this analysis, effects of sample location could be determined, but were weak. Importantly, the dominant part of the pine pollen microbiome (i.e. the fungi), exhibited stability or conservation in community structure irrespective of sampling location. No change in geographic-related sampling was evident. Rather, it was components of the bacterial community in which changes in the abundance of some taxa occurred in association with sampling distance.

Clostridia was the most notable taxa to change in abundance over space, exhibiting a roughly west-to-east gradient in distribution (from high to low abundance, respectively). Sphingobacteriia was the most abundant species and was present in nearly all samples, whereas Acidimicrobiia, Deinococci and Oligoflexia were only present in 1-3 samples with no obvious correlation. In these instances, we hypothesise that environmental filtering of specific bacterial taxa may be occurring, where perhaps factors related to tree health or other factors within the environment are influencing colonisation onto tree tissues. We observe the legacy of this within the pollen microbiome simply as a surface where microbes from the environment can be deposited.

Likewise, there were no significant differences in fungal and bacterial community composition across the two annual time points. However, with regards to bacteria, some distinctions emerged. In 2020, certain classes such as Verrucomicrobiae, Thermomicrobia, Phycisphaerae, Oligoflexia, Acidiimicrobia, and Acidobacteria Groups 3, 6, and 7 were exclusively detected, while classes including Methanobacteria, Gemmatimonadetes, and Bacteroidia were only identified in 2021. Some changes to the microbiome are expected, whether arising from random fluctuations or variations in sampling from one year to the next. It is notable, however, that like the results for the geographic distance testing, that changes over years were observed in the bacterial and not fungal component of the microbiome. This provides additional support for the bacterial microbiome being of secondary significance to the overall pollen microbiome; its assembly isn’t as tightly regulated or consistent as that for fungi and are more stochastically influenced by environmental site or sampling time. It’s important to note that the sampling methods remained consistent between the two time points, and the timing of the sample collection was the same. However, factors like the pollen release from catkins, seasonal changes and weather conditions are all variable. Overall, it is evident that neither sampling year nor geographic distance are the main dominant drivers of the pine pollen microbiome, particularly the dominant fungal community component which demonstrated remarkable stability.

### Is the microsporangiate a key source of the pollen microbiome?

Examination of the *P. radiata* catkins and pollen by SEM revealed that catkins had visibly rich microbial growth on its surface, unlike the pollen grains which showed relatively sparse microbial presence (Suppl. Fig S4). Pollen is formed within, and released from, the male microsporangiate strobili. Microsporangia within the pollen cones produce microspores through meiosis and these develop into pollen grains. As such, the formation of pollen occurs within the enclosed cones, being isolated from the external environment until the cones mature and open for pollen release. Accordingly, the developmental opportunity for external colonisation of pollen with microbiome is limited; microbiomes present on pollen are likely to be sourced from the surrounding catkin material. It is likely, therefore, that movement of microbiomes from the catkin to the pollen is an important primary source of the pollen microbiome. Indeed, there has been a long history of use of bleach treatment of catkins to reduce mould and other growth on pine pollen (Tulecke, 1954), further demonstrating fungi present on pollen originates from surface of the catkins. If validated, host influence of the pollen microbiome may occur via properties related to the chemistry and ecology of the catkin structure itself, and focus on the metabolome and ecology of this tissue may shed light on processes affecting assembly and transmission of the pollen microbiome.

Not all the microbiome may reside on the surface of pollen. Following sterilisation, growth of endophytic microbiota, such as *Enterobacter cloacae*, has been demonstrated in a range of *Pinus* spp (Madmony *et al.,* 2005). The functional significance of these isolates is unknown, however strong co-evolutionary processes typically drive plant-endophytic associations (Lyu *et al.,* 2012; Koskella and Bergelson, 2020) and, as such, shared fitness outcomes are likely for both the plant and microorganism.

### Pine pollen has a core microbiome

There was a consistent presence of specific fungal and bacterial taxa across *P. radiata* pollen samples; this indicates that pine pollen holds symbiosis with a core microbiome. This core microbiome is, axiomatically, the stable fraction being consistently present (see below) over two years of sampling and across locations spanning broad geographic range and proximity to land uses. Indeed, the term ‘core microbiome’ is described as one that hosts anywhere from 30-95% of microbial taxa (Risely, 2020) across samples, depending on the habitat being investigated. In our study, we define a core member as having a 50% prevalence and 0.01% relative abundance detection threshold (Graystock *et al.,* 2017; Kardas *et al.,* 2023). From these parameters, five bacterial and ten fungal core genera were identified. At a greater taxonomic resolution such as phyla, the dominant bacterial and fungal phyla on radiata pollen was Proteobacteria and Ascomycota, which are also dominant on pollen for a range of wind and insect pollinating plants (Manirajan *et al.,* 2018). At the genera level, *Robbsia, Sphingomonas, Hymenobacter, Acidisoma* and *Methylobacterium* were the dominant genera of the core bacterium community. Of these, *Robbsia* and *Methylobacterium* have been found on birch and canola pollen previously (Shi *et al.,* 2023; Manirajan *et al.,* 2018), and *Sphingomonas* are well documented as a core bacterium via in pollen collected by bees (Dew *et al.,* 2020). *Methylobacterium* are a genus of well-documented phyllosphere epiphytes that secrete cytokinins and auxin plant hormones (Sanjenbam *et al.,* 2022). *Hymenobacter* and *Acidisoma* have not yet been reported to be associated with pollen, but are known plant phyllosphere bacteria (Ares *et al.,* 2021; Reis *et al.,* 2015).

The core fungal genera were *Genolevuria, Microsphaeropsis, Gelidatrema, Epicoccum, Perusta, Taphrina, Vishniacozyma, Hormonema, Naganishia*, and *Symmetrospora*. The fungal families Didymosphaeriaceae and Dothioraceae, Taphrinaceae were also found in the 8 pollen species studied by Manirajan *et al* 2018, with Dothioraceae being prominent throughout most of the pollen species. This aligns with our study of pine pollen too, where Dothideomycetes was the most abundant class of fungi present.

The presence of Dothideomycetes is not surprising as it contains a wide range of plant-associated fungi with ecological roles ranging from saprophytes, pathogens, to endophytes (Schoch and Grube, 2007). It is also the dominant fungal group on *P. radiata* needles (Addison *et al.,* 2023). Not surprisingly there were more fungal core groups compared to bacterial groups, which is expected due to the sheer number of fungal groups recorded compared to bacterial groups. These results support those of Manirajan *et al* (2018) who report more core fungi than bacteria on pollen from eight plant species, despite these pollens all having greater bacterial richness.

Comparing our results to the core microbiomes from the species of pollen Manirajan *et al* 2018 studied, of which there were 33 fungal core genera and 12 bacterial core genera, only Vishniacozyma and Taphrina fungi were shared with these other pollen species. This shows that the pollen microbiome is indeed species specific, giving rise to unique groups especially at finer taxonomic resolution such as genera. As before, this provides support that the host plant is key in supplying/recruiting or maintaining specific and stable microbiome community. At greater taxonomic resolution such as phyla, taxonomic lineages of both bacteria and fungi in this study (i.e. dominance of Proteobacteria and Ascomycota) appear to be consistent across other pollen microbiome studies.

### Limitations and future research

Due to the experimental system used, this study was conducted on field-collected pollen specimens from *P. radiata*. Control of tree genotype was not conducted, nor age of trees measured. However, all were likely originated from commercial stock of *P. radiata* domesticated from parent material originally sourced from California via Australia (Burdon *et al.,* 2017). Given the findings reaffirm the seminal role of the host (as opposed to the environment) in shaping pollen microbiome communities, further work should consider inclusion of host genotype in their design. Further to this, locations in which trees were sampled ranged from road-side shelterbelts through, small stands, and single trees in recreational parks. These factors were not (and could not be) formally tested within the analysis, yet possibly contribute to some of the total variation captured in the ‘environmental’ testing. However, as this was found to be of limited importance in microbiome assembly anyhow, we propose that further investigation of this may not be needed (e.g. structured sampling of pines based on adjacent land use) for the pine microbiome. Finally, only visibly heathy trees were chosen for this study. Thus, influence of stress from factors such as drought, disease, pollution etc could not be assessed, but may be important (Obersteiner *et al.,* 2016). These should be considered in future research.

The domestication and economic importance of the species means that a wealth of information continues to be generated on its wider biology, including physiology and genetics. Robust systems are in place for propagation, transformation and other experimentation on *Pinus radiata*, and a wide network of existing trials are in place to test aspects of the tree’s phenotype and fitness in diverse environments (Burdon *et al.,* 2017). Similarly, an increasing wealth of information is being built on the microbiome of *P. radiata* including needles, roots, wood, and rhizosphere (Addison *et al.,* 2023a,b; Prihatini *et al.,* 2015; Rúa *et al.,* 2016). The results presented here have shown the importance of considering the pollen microbiome when examining the overall microbiome of trees. Further research investigating whether this microbiome is passed down vertically to seeds and seedlings will be important. Moreover, exploring whether the influx of microbes each spring plays a role in shaping the phyllosphere, litter layer, or soil microbiomes, as well as in the dissemination of pathogens. This work will go towards answering key questions related to the assembly and function of the *P. radiata* tree-microbiome system, improving the fitness and resilience of forests in a rapidly changing world.

## Acknowledgments

This work was supported by the Ministry of Business, Innovation and Employment within ‘The Tree Microbiome Project: at the root of climate proofing forests (C04 × 2002)’, New Zealand’s Forest Growers Levy Trust, and via Strategic Science Investment Fund allocation to Scion within the ‘Resilient Forests’ research programme. The authors thank Laureline Rossignaud for assistance in sampling in 2021. SEM work was conducted at University of Canterbury Engineering Department with guidance by Shaun Mucalo.

## Data Availability

Sequencing data from this project can be found on the Short Read Archive (SRA) database, under the BioProject accession number PRJNA1109979

