## Supplementary Figures for "Pollen Partners: The Symbiotic Microbes of *Pinus radiata* Pollen"

---

---

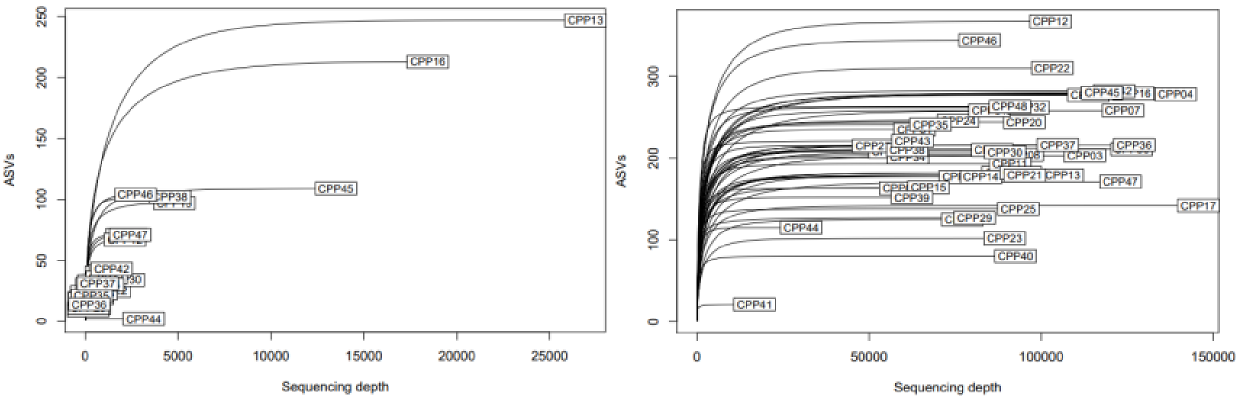


**Figure S1**: Rarefaction curves for bacteria (left) and fungi (right) MiSeq-based sequencing of *Pinus radiata* pollen samples. Each curve is the result of sequencing of an individual sample of pollen collected from a single location and single time point.

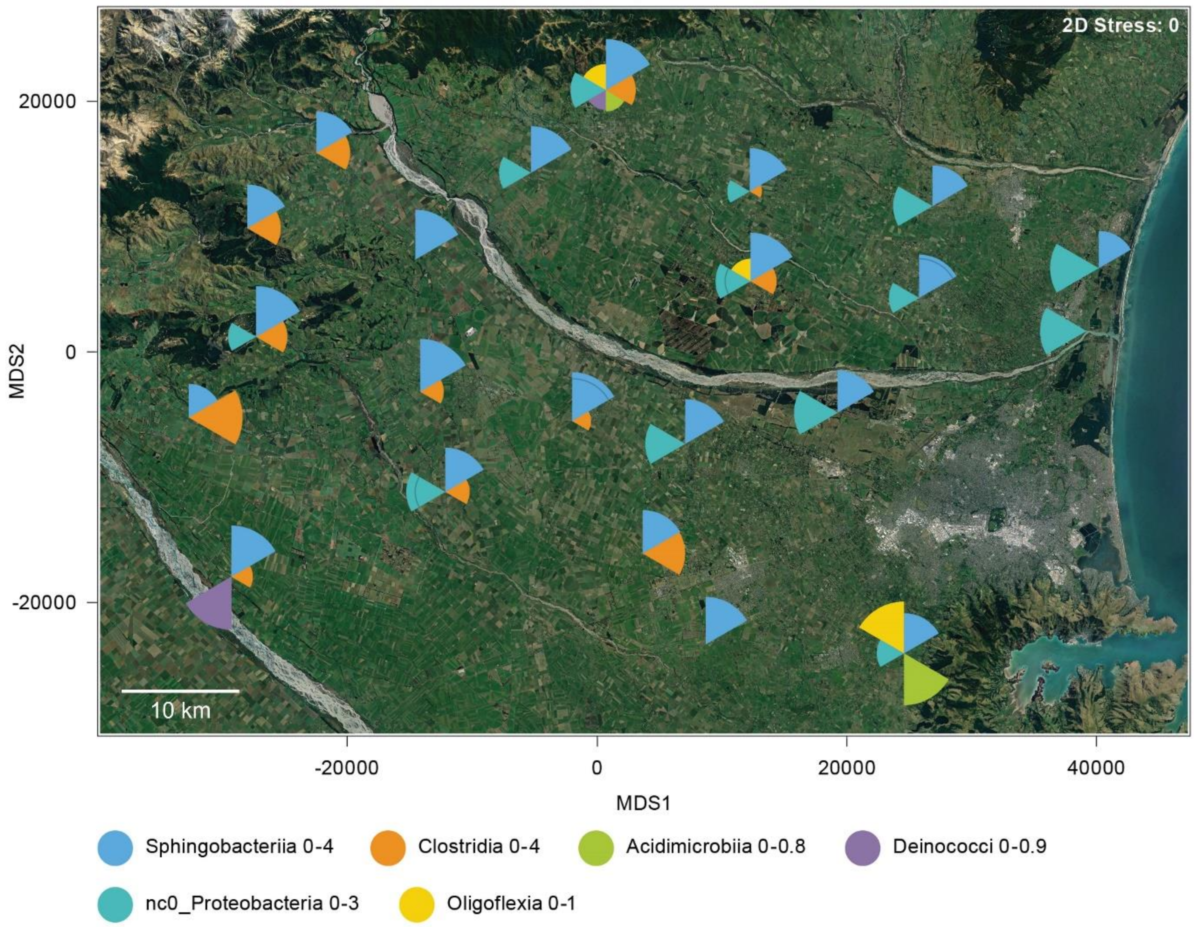


**Figure S2.** Overlay plot showing changes in abundances of bacterial groups found to have associations with geographic distance (BIO-ENV testing). The plot is superimposed over the Google Earth satellite imagery of the sampling area. Bacterial abundances are on an assorted scale (see key); these values are derived from the abundance-standardised, square-root transformed Class aggregated values. The scale (derived through square root transformation of relative abundances) shows the range of classes present across samples. Each segment is reflective of the square-root transformed relative abundance with segment size proportional to the scale shown. Sizing of segments are analogous to a relative abundance of 0-16%,0-9%,0-1%, 0-0.81% and 0-0.64% when back-transformed from scale shown in figure i.e. 0-4, 0-3,0-1, 0-0.9 and 0-0.8 respectively. Fungal taxa showed no evidence for association between changes in abundances and geographic range and, therefore, are not shown.

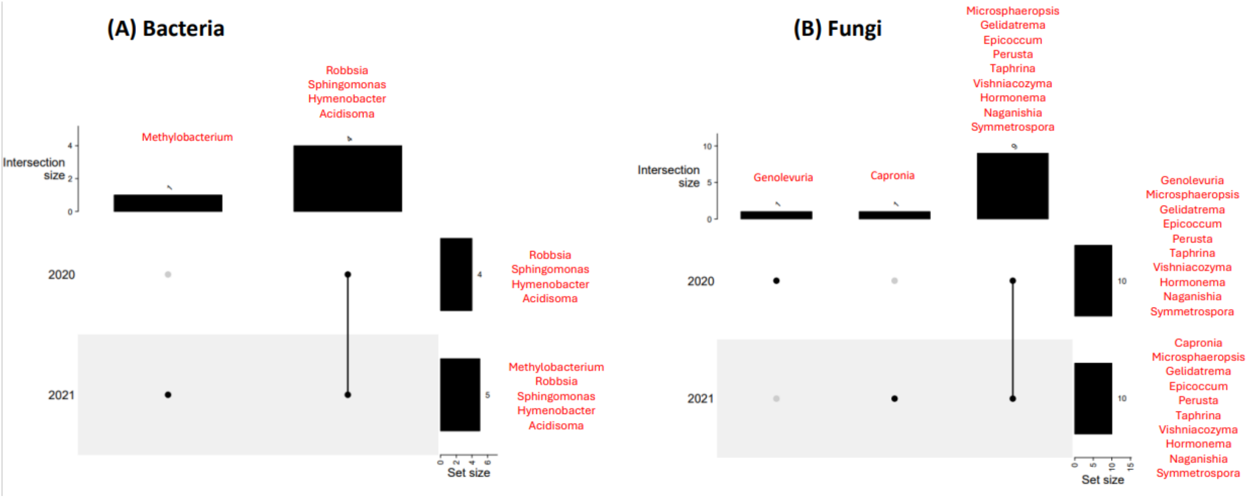


**Figure S3**: Core genera of bacteria (left) and fungi (right) between and across years. Solid black dots indicate presence, grey dots indicate absence, lines connecting solid black dots means presence shared across years. Vertical bars are total number of core genus either between (i.e. if black dots are connected with lines)/within years (i.e black dots without connecting lines on figure) while horizontal bars are total within each year

Year-based core genera (single black dots on figure) of bacteria and fungi are given in Figure S3 A and S3 B, respectively. These were determined using a 50% prevalence and 0.01 detection threshold. For year-specific effects, the bacterial genus Methylobacterium (Alphaproteobacteria) exhibited collection date effects, being core across samples collected in 2021 only (Fig S3A). For the fungi, Genolevuria (Basidiomycete) was core in 2020 only, and Capronia (Ascomycete) in 2021 only (Fig. S3B). Apart from this single variation, 4 and 9 core genera were found to be shared- (black lines connecting dots from 2020 and 2021 with vertical bar indicating counts shared between year 2020 and 2021) between years for bacteria and fungi respectively. Within each year (i.e. horizontal bars), fungal groups recorded 10 core genera with 9 of these being constant irrespective of year (i.e. Miscrosphaeropsis, Gelidatrema, Epicoccum, Perusta, Taphrina, Visniacozyma, Hormonema, Naganisha, Symmetrospora). For bacterial groups, 4 core genera were found in 2020 (i.e. Robbsia, Sphingomonas, Hymenobacter, Acidisoma) while 5 were found in 2021 (i.e. Methylobacterium, Robbsia, Sphingomonas, Hymenobacter, Acidisoma

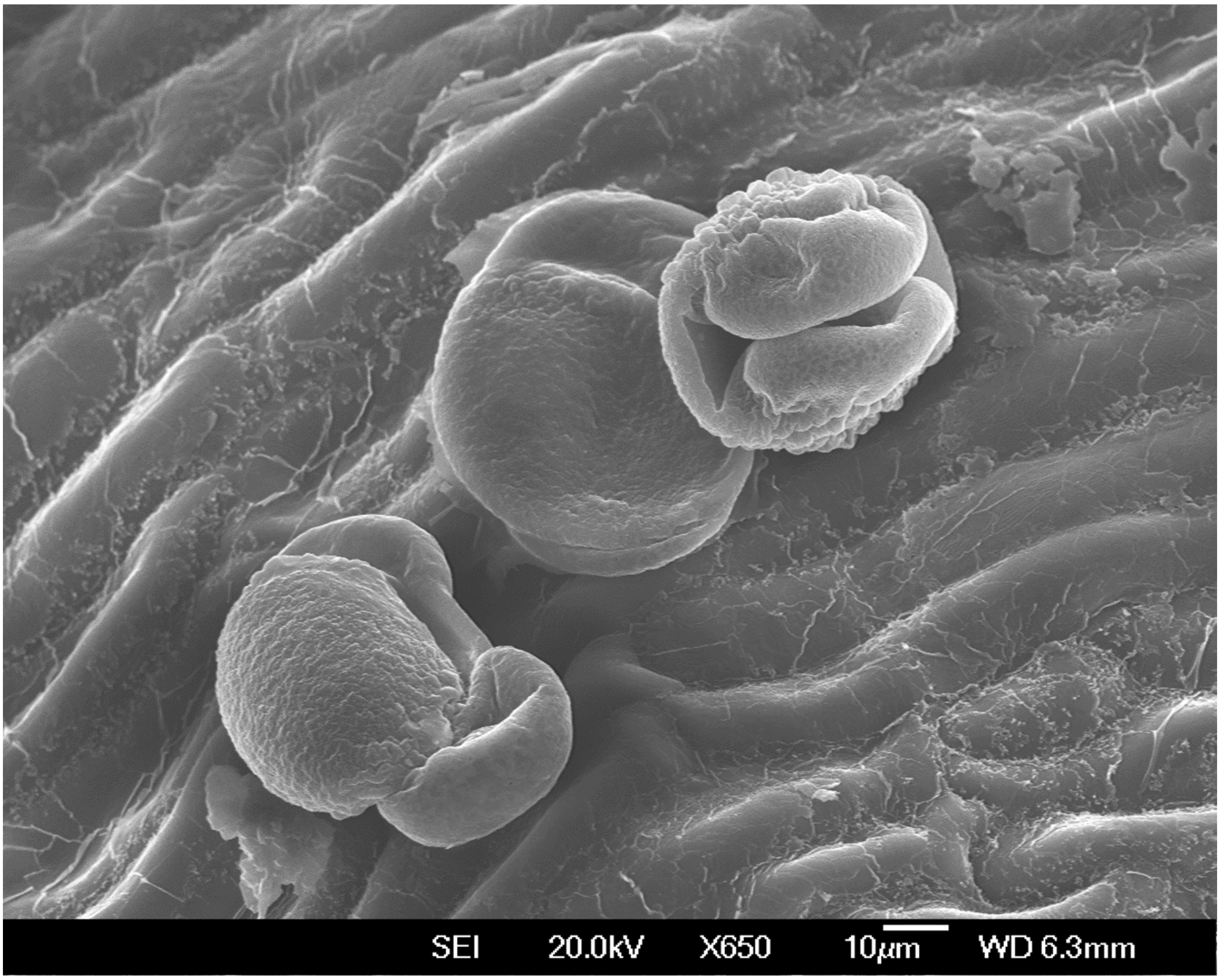


**Figure S4:** Scanning electron microscope (SEM) image of *P. radiata* pollen grains on microsporangiate strobili surface; scale bar 10 µm.
